# Vimentin regulates mitochondrial ROS production and inflammatory responses of neutrophils

**DOI:** 10.1101/2024.04.11.589146

**Authors:** Thao Ngoc Huynh, Jody Toperzer, Allison Scherer, Anne Gumina, Tonya Brunetti, Michael K. Mansour, David M. Markovitz, Brian C. Russo

## Abstract

The intermediate filament vimentin is present in immune cells and is implicated in proinflammatory immune responses. Whether and how it supports antimicrobial activities of neutrophils is not well established. Here, we developed an immortalized neutrophil model to examine the requirement of vimentin. We demonstrate that vimentin restricts the production of proinflammatory cytokines and reactive oxygen species (ROS), but enhances phagocytosis and swarming. We observe that vimentin is dispensable for neutrophil extracellular trap (NET) formation, degranulation, and inflammasome activation. Moreover, gene expression analysis demonstrated that the presence of vimentin was associated with changes in expression of multiple genes required for mitochondrial function and ROS overproduction. Treatment of wild-type cells with rotenone, an inhibitor for complex I of the electron transport chain, increases the ROS levels. Likewise, treatment with mitoTEMPO, a SOD mimetic, rescues the ROS production in cells lacking vimentin. Together, these data show vimentin regulates neutrophil antimicrobial functions and alters ROS levels through regulation of mitochondrial activity.

## Introduction

Intermediate filaments are ubiquitous cytoskeletal proteins in mammalian cells. There are greater than 70 different intermediate filament proteins in humans (1–4). Monomers of intermediate filament proteins can homodimerize or heterodimerize depending on the type, and they can assemble into fibers that lack polarity (1). Networks of intermediate filament fibers provide structural support to cells and are highly dynamic (5, 6). Intermediate filaments are important for positioning organelles and for cell motility, among other activities (7–10). They are required for the virulence of many bacterial pathogens and their loss is correlated with inflammatory bowel disease severity (11–13). Moreover, they are associated with cancer progression, neuropathies, and acute lung injury (14–16). Thus, intermediate fliaments provide basic functions required for cellular homeostasis, and their dysregulation occurs in a variety of diseases associated with significant morbidity in humans.

Vimentin, the most abundant intermediate filament protein (17), is thought to be important for regulating inflammatory response during infection for several reasons: 1. Vimentin is expressed in immune cells, including macrophages and neutrophils (18, 19); 2. In macrophages, vimentin regulates the NADPH oxidase complex and functions as a scaffold for NLRP3 inflammasome activation (14, 15); 3. Whereas the loss of vimentin in mice leads to no overt phenotype at baseline (20), a thorough characterization of *Vim^-/-^*mice under stress conditions reveals prominent abnormalities (21, 22); 4. Recent studies showed that mice with vimentin are more susceptible to colitis compared to *Vim^-/-^* mice, likely due to its interactions between vimentin and the p47phox active subunit of NADPH oxidase (15). These data together suggest that vimentin may be generally required for immunological responses to pathogens. Whereas the role of vimentin in macrophages has been studied, how vimentin functions in neutrophils has been largely overlooked.

In this study, we demonstrate that vimentin affects the function of neutrophils by regulating mitochondrial reactive oxygen species (ROS) production. By using immortalized neutrophil progenitor cells isolated from wild-type (*Vim^+/+^*) and *Vim^-/-^* mice, we showed that vimentin represses the levels of ROS production by regulating the mitochondria. Moreover, vimentin presence was associated with neutrophil swarming and phagocytosis, but its loss increased TNFα and IL-6 cytokine production. Interestingly, vimentin was not required for inflammasome activation, demonstrating a distinct role for vimentin in the neutrophil inflammatory response as compared to its function in macrophages (14). Taken together, we suggest a model in which vimentin has diverse roles in the inflammatory response of neutrophils.

## Materials and Methods

### Generation of HoxB8-immortalized neutrophil progenitor cells and differentiation of neutrophils

Bone marrow was isolated from wildtype and vimentin knockout mice at the University of Michigan (18). Bone marrow was stimulated with recombinant murine IL-3 (10 ng/mL) and IL-6 (10 ng/mL) (Peprotech, 21616) for 48 hours. Cells were then transduced with ER-HoxB8 lentivirus (23) and were cultured in RPMI (Fisher, 32404014) containing 10% FBS (Biotechne, S11150H), 5% stem cell factor (SCF) (described below), and 10 μg/mL β-esterdiol (Sigma, E2758-1G). Following G418 selection (2 mg/mL), individual cells were sorted and clonally expanded in media to maintain cells in the progenitor state. 5_X_10^5^ cells were collected and differentiated as previously described (24).

SCF was generated by culturing CHO cells in IMDM (ThermoFisher, 12440061) supplemented with 10% FBS (Biotechne, S11150H). Cells were cultured for 48 hours beyond confluency. The media was then collected and replaced every 24 hours for 72 hours. The collected media was pooled and used as conditioned media for SCF above.

### Flow Cytometry

The Fc receptors were blocked on 5_X_10^5^ neutrophils resuspended in PBS with 5% FBS and TruStain FcX (1:50, BioLegend, 101320). Neutrophils were incubated with FITC conjugated anti-mouse Ly6G (1:50, BD Biosciences, 551460) and PE conjugated anti-mouse/human CD11b (1:200, Biolegend, 101207) for 30 minutes on ice. Samples were washed with FACS buffer, and fixed with FluoroFix (Biolegend, 422101) for 20 minutes on ice. Samples were resuspended in FACS buffer and analyzed using a BD LSRFortessa X-20.

### Western blots

Western blots were incubated with anti-vimentin (1:10000, Biolegend, 919101) or anti-caspase-1 (1:100, Abcam, ab207802) antibodies overnight at 4°C, washed with TBS-T, and incubated with HRP-conjugated secondary antibodies [anti-chicken (1:5000, Jackson Immuno Research, 103-03-155) or goat anti-rabbit] for two hours at room temperature. β-actin (Sigma, A3845) was used as a loading control. Western blots were developed on a Syngene G:BOX chemi XX6 using West Pico (Thermo Scientific, 34580). Band intensity was quantified using ImageJ.

### Nuclear staining

For each clone tested, 5_X_10^5^ neutrophils were incubated with cell tracker red (Invitrogen, C34552) for 30 minutes at 37°C. Cells were washed with media and transferred to an 18-well chamber slide (ibidi 81817). Hoechst (Invitrogen, H3570) was added to the media, and the cells were incubated for 30 minutes prior to imaging.

### Swarming assay

*C. albicans* producing the fluorescent protein iRFP670 was cultured at 30°C in YPD media containing nourseothricin (Jena Bioscience, AB-101-50ML) (25). On the day of experiment, *C. albicans* was diluted 1:1000 in YPD containing nourseothricin and 10% FBS and incubated for 2 hours at 30°C to promote hyphae formation. *C. albicans* was added to wells of a Proplate (Grace Biolabs, 204860) that was micro-arrayed, as previously described (26), to enable *C. albicans* to adhere to the surface by centrifugation at 800 _X_*g* for 10 minutes. Unadhered *C. albicans* was removed by washing thoroughly with PBS. 1_X_10^5^ neutrophils were stained with cell tracker green (Invitrogen, C2925) and added to each well. Microscopy-based images were collected such that tiled images examined 32 microspotted positions per well. To quantify the efficiency of neutrophil swarming, green fluorescence from neutrophils recruited to each microspot containing *C. albicans* was measured.

### ELISA

Neutrophils were seeded at 1_X_10^5^ cells per well in a 96-well flat-bottom plate in 250 µL of media. Cells were stimulated with LPS (Invivogen, tlrl-3pelps) at 25 ng/mL and incubated at 37°C with 5% CO_2_ as indicated for each experiment. The supernatant was collected at indicated times and protein concentrations were determined by ELISA following the manufacturer’s instructions for IL-6 (R&D systems, DY406-05), TNFα (R&D systems, DY410-05), Matrix metalloprotease 9 (MMP-9) (R&D Systems, DY6718), and IL-1β (R&D Systems, Y401-05).

### Quantification of inflammasome activation

Neutrophils were seeded at 1_X_10^5^ cells in a 96-well flat-bottom plate in 150 µL of media. For stimulated samples, LPS (Invivogen, tlrl-3pelps) was added at a final concentration of 1.67 µg/mL. The neutrophils were incubated for 1.5 hours at 37°C with 5% CO_2_, then 100 µL of media containing 7.4 µg/mL nigericin (Invivogen, tlrl-nig) was added. The cells were incubated for an additional 40 minutes. Then, 50 µl of supernatant was collected every two hours and used to quantify cytokine abundance by ELISA.

### Neutrophil extracellular trap formation

7.5_X_10^4^ neutrophils were seeded in an 18-well chamber slide coated with fibronectin (Sigma, F1141). Cells were seeded in RPMI with 0 µM or 1.25 µM calcium ionophore A23187 (Sigma, C7522), and incubated at 37°C with 5% CO_2_ for 1.5 hour. Cells were washed with PBS and imaged in the presence of Hoechst and sytox green.

### Phagocytosis

2.5_X_10^4^ neutrophils were seeded into an 18-well chamber slide coated with fibronectin. *E. coli* pBR322-GFP were added at a multiplicity of 20 and centrifuged at 800 _X_*g* for 10 minutes. Following 30 minutes of incubation at 37°C, the neutrophils were washed eight times with media and fixed with paraformaldehyde. Extracellular *E. coli* were determined by staining with rabbit anti-*E.coli* (1:100, Abcam, ab137967) for 1 hour at 25°C and then goat anti-rabbit antibody conjugated with Alexa Fluor 750 (1:100, Invitrogen, A21039) for 2 hours at 25°C. Nuclei were stained with Hoechst (1:500; Invitrogen, H3570). Intracellular bacteria per neutrophil were quantified.

### Quantification of ROS

Reactivity of 1_X_10^5^ neutrophils in HBSS containing 100 µM cytochrome c (Sigma, C7752) was determined by absorbance at 550 nm and 490 nm every 2 minutes for 2 hours using a Biotek synergy H1 (24). Neutrophils were stimulated with 100 nM PMA (Sigma, P8139). For studies investigating whether mitochondrial function altered ROS production, neutrophils were treated with 5 μM rotenone or 250/500/1000 μM mitoTEMPO in HBSS.

### RNA extraction and sequencing

Neutrophils were treated with 25 ng/mL LPS (Invivogen-tlrl-3pelps), or left untreated, at 37°C with 5% CO_2_ for 1 hour. RNA was extracted using the RNeasy micro kit (Qiagen, 74004) and DNase treatment occurred using Turbo DNA-free DNAse (Qiagen, AM2238). Samples were prepared using the Universal Plus mRNA-Seq library preparation kit with NuQuant and sequenced on an Illumina NovaSeq 6000 with 2x150 bp sequencing at the University of Colorado Anshutz Medical Campus Genomics Core. Quality of reads was assessed pre- and post-trimming using FastQC v0.11.9 (27). Illumina universal adapters were removed, bases were trimmed if the Phred Score was less than 24, and after trimming, any reads that were fewer than 20 base pairs in length were discarded using Cutadapt v4.4 (28) under Python version 3.10. Reads were aligned and quantified to the mm10 reference genome using STAR v2.7.10b (29). The quality of the alignments were assessed using various tools including samtools v.1.17 (30), PicardTools CollectRnaSeqMetrics v2.27.5-13-g04e9b2 (31), and the summarization of log outputs generated by STAR. Differential gene expression was assessed using two R packages: RUVseq v1.20.0 (32) and DESeq2 v1.26.0 (33). RUVseq was used to identify covariates to address any batches and sources of unwanted variation to regress from the normalized gene expression matrix using a GLM. We identified four RUV covariates to add to the model and subsequently used DESeq2 to normalize and statistically run the GLM for differential expression analysis. fGSEA was performed using the R package fgsea v1.22.0 (34), and ranks were based upon the Wald statistic extracted from the differential expression matrix generated by DESeq2. The Hallmark gene set from the Molecular Signatures Database (MSigDB) was used for enrichment analysis, and pathways were considered statistically significant if they had an adjusted p<0.05. Venn diagrams of pathways were based upon the leading edge genes identified in the fGSEA.

### Quantification of mitochondria and mitochondrial ROS

Cells were resuspended in HBSS containing 10% FBS, 200 nM mitosox red, and 40 nM mitotracker green and incubated at 37°C with 5% CO_2_ for 30 minutes. Cells were washed twice with HBSS and imaged. Images were deconvolved and intensity per cell was determined for mitosox and mitotracker.

### Microscopy

Fluorescent images were acquired on a Nikon Eclipse Ti-2 inverted light microscope equipped with an Orca Fusion BT cMOS camera (Hammamatsu), an IRIS 15 cMOS camera (Photometrics), Semrock Brightline filters, and an Oco stage top incubator. NIS elements software (Nikon) was used for image acquisition and analysis. Images were deconvolved using a Richardson-Lucy algorithm with 20 iterations. Microscopic images were pseudo-colored and assembled using Adobe photoshop or FIJI (NIH). Images were collected randomly across the coverslip using an automated imaging pipeline created in NIS elements, or the samples were blinded prior to imaging.

### Statistics

Statistical difference between means was analyzed with GraphPad Prism 9. Statistical difference of two means was determined either by Student’s t-test or a paired t-test. Unless indicated otherwise, the differences between the means of multiple groups were determined either by two-way ANOVA with Sidak’s *post hoc* test or by one-way ANOVA with Holm-Sidak’s *post hoc* test.

## Results

### Generation of neutrophils from immortalized stem cell lines

Vimentin is required for macrophages to respond to pathogens (14, 18, 35), but its requirement for neutrophil function is uncertain. To investigate the role of vimentin in neutrophils, we generated immortalized neutrophil progenitors from the bone marrow of wild-type (*Vim^+/+^*) and vimentin knock-out (*Vim^-/-^)* mice by transduction of the transcription factor HoxB8. We isolated five or six clonal lines of granulocyte monocyte progenitors (GMPs) from the bone marrow of wildtype (*Vim^+/+^*) and vimentin knockout (*Vim^-/-^*) mice, respectively, and confirmed the expression or lack of vimentin in these cells by western blot (Figure 1A-B). The activity of HoxB8 was regulated by fusion to the estrogen receptor binding domain, such that HoxB8 is only active in the presence of estrogen. The removal of estrogen from the culture media and the addition of IL-3 and granulocyte colony-stimulating factor (GCSF) enabled differentiation of the GMP into mature neutrophils, which was evaluated by cell surface expression of CD11b and Ly6G (Figure 1C-D) and by examining nuclear morphology (Figure 1E-F). Whereas the GMPs displayed round nuclei and were CD11b low and Ly6G negative (Figure 1C-F), differentiated neutrophils displayed lobular nuclei and were CD11b positive and Ly6G positive (Figure 1C-F). Thus, neutrophils efficiently differentiate from progentitors derived from mice that produce or lack vimentin.

**Figure 1.**
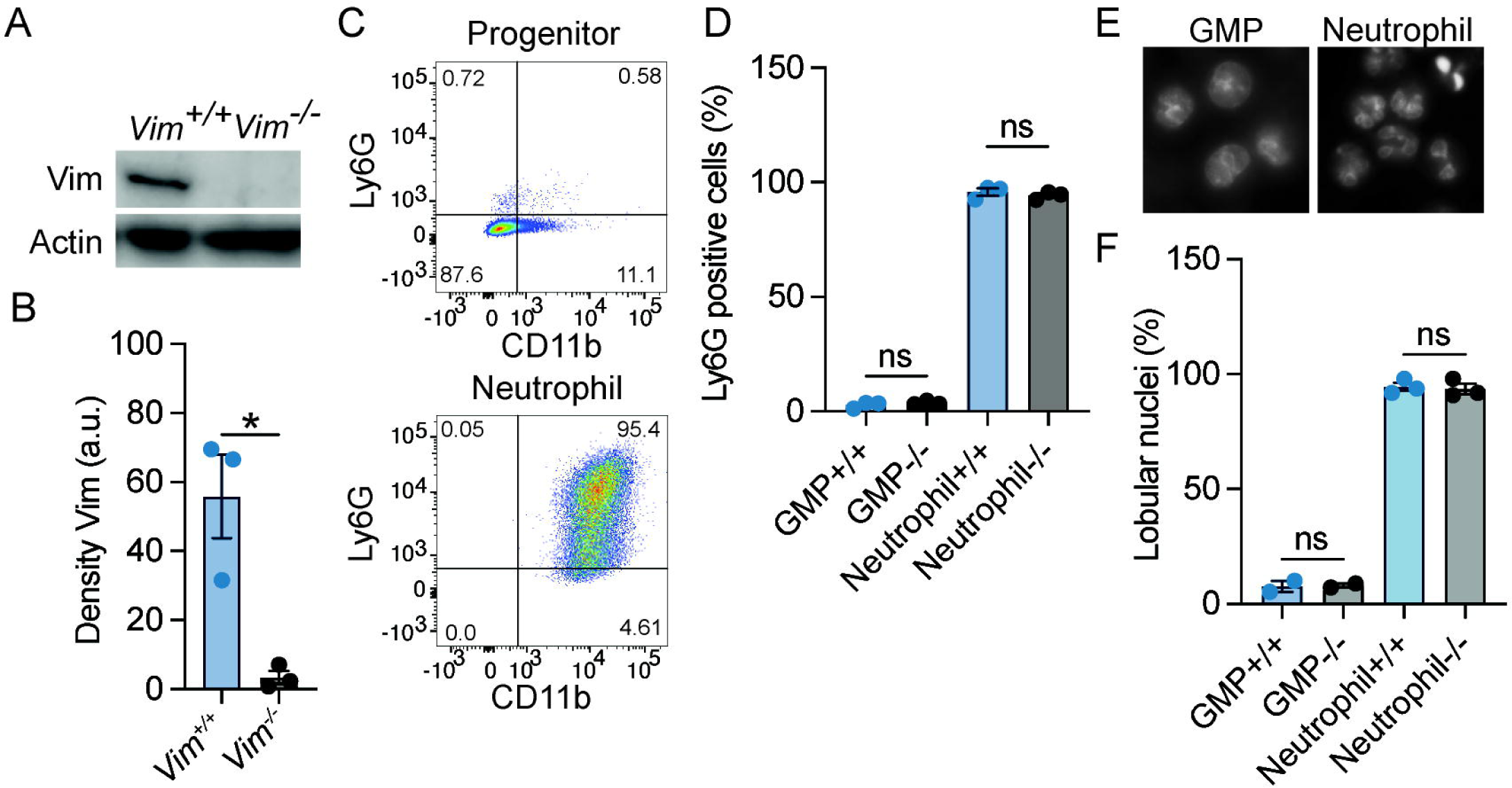
HoxB8-transformed wild-type and *Vim^-/-^* GMP differentiated into neutrophils. A) Representative image of western blot showing vimentin expression. B) Quantification of the western blots represented in panel A. C) Ly6G and CD11b expression for neutrophil progenitor cells and differentiated neutrophils. D) Quantification of the percentage of Ly6G positivity. E) Representative images of the morphology of the nuclei of GMP and neutrophils. F) Quantification of percentage nuclei that are globular. (B, D, and F) Dots are independent experiments using 5 *Vim^+/+^* and 6 *Vim^-/-^*cell lines. Data are mean ± SEM. *: p < 0.05; ns, not significant; paired t-test (B) or one-way ANOVA with Sidak’s *post hoc* test (D and F).

### Vimentin is required for swarming and phagocytosis, but not NETs or degranulation

Neutrophils are the most abundant white blood cell in the body and are key participants in the defense against fungal and bacterial infections (36, 37). In some cellular systems, the presence of vimentin is correlated with the efficiency of cell motility (2, 8, 38–42), suggesting that the ability of neutrophils to swarm to pathogens could be impacted by the loss of vimentin. We tested the ability of *Vim^+/^*^+^ and *Vim^-/^*^-^ neutrophils to swarm. We observed that, at 4 hours of co-incubation, there was a 45.3% reduction in the efficiency of *Vim^-/-^* neutrophils to swarm to *C. albicans* as compared to *Vim^+/^*^+^ neutrophils (Figure 2A-B). Thus, vimentin enhances the ability of neutrophils to swarm.

**Figure 2.**
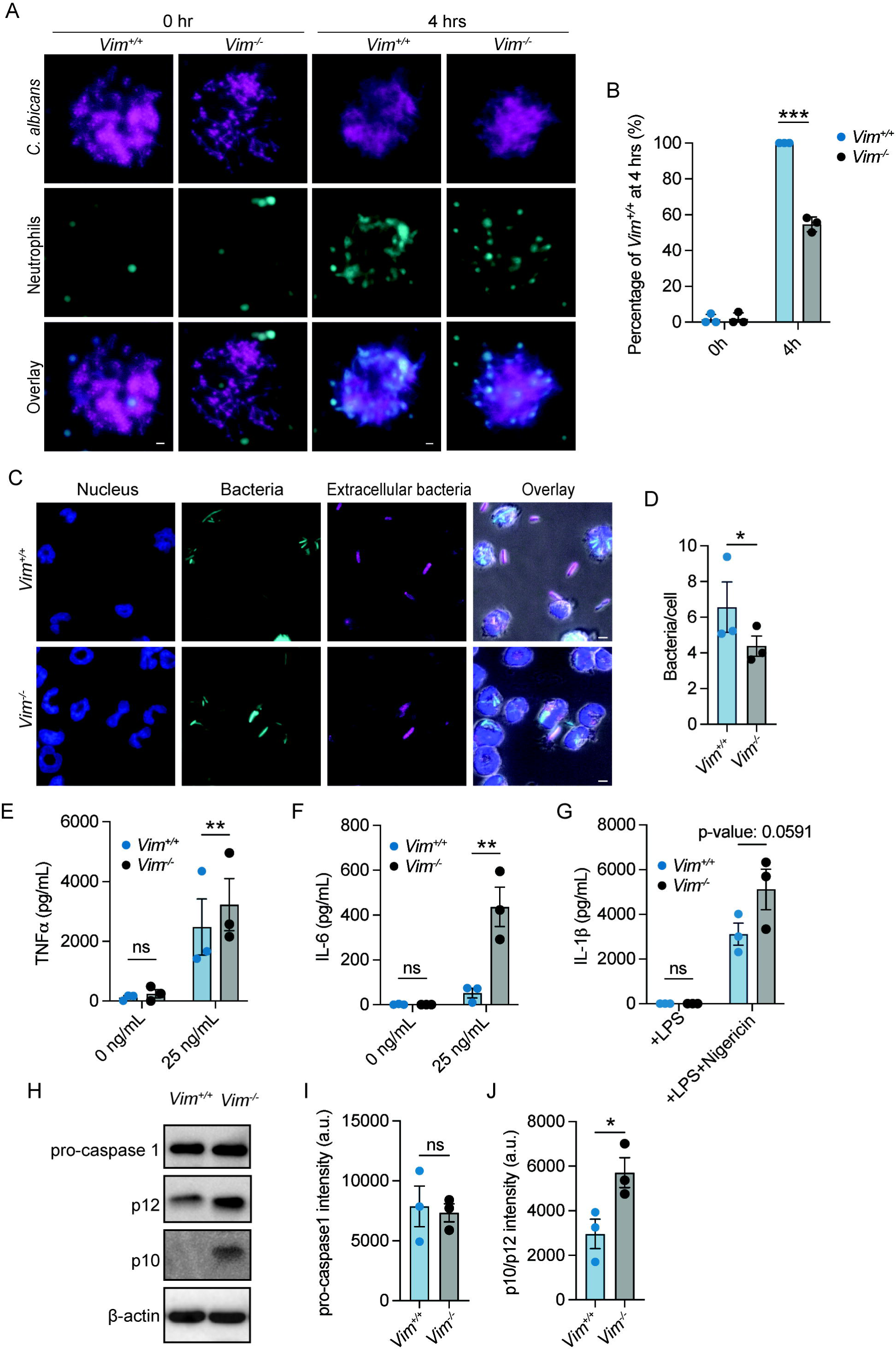
Functional analysis of neutrophils with and without vimentin. A) Representative images of *Vim^+/+^* and *Vim^-/-^* neutrophils swarming to C. *albicans.* Cyan, neutrophils; magenta, *C. albicans*. Scale bar: 10 μm. B) Quantification of neutrophil swarming efficiency by intensity. Swarming of *Vim^+/+^* neutrophils is 100%. C) Representative images of phagocytosis of *E. coli* by *Vim^+/+^*and *Vim^-/-^* neutrophils. blue, DNA; cyan, all *E. coli*; magenta, extracellular *E. coli*. Scale bar: 5 μm. D) Quantification of number of intracellular bacteria per cell in *Vim^+/+^* and *Vim^-/-^*neutrophils. (E-F) Quantification of TNFα (E) or IL-6 (F) produced by *Vim^+/+^* and *Vim^-/-^* neutrophils in response to LPS stimulation at 6 hours. (G) Quantification of IL-1β produced by *Vim^+/+^* and *Vim^-/-^* neutrophils in response to treatment with LPS alone or LPS and nigericin at 2 hours. (H) Representative western blot for pro-caspase-1, p12, and p10 after LPS and nigericin (to induce inflammasome activation) stimulation of *Vim^+/+^*and *Vim^-/-^* neutrophils. (I-J) Quantification of pro-caspase-1 (I) and p12/p10 levels (J). (B, D, E-G, and I-J) Data mean ± SEM. Dots are independent experiments. ns, not-significant; *, p <0.05; **, p<0.01; ***, p<0.001 by two-way ANOVA with Sidak’s *post hoc* test (B, E-G) and paired t-test (D, I, and J).

In addition to swarming to the site of pathogens, neutrophils phagocytose and kill pathogens (37). The role of vimentin in phagocytosis was determined by examining the efficiency by which *Vim^+/^*^+^ and *Vim^-/^*^-^ neutrophils phagocytosed *E. coli. Vim^-/-^* neutrophils showed fewer intracellular bacteria (∼4.3 bacteria/cell) compared to *Vim^+/^*^+^ neutrophils (∼6.5 bacteria/cell) (Figure 2C-D). Although, there was no difference in the number of *Vim^+/^*^+^ or *Vim*^-/-^ neutrophils that could phagocytose *E. coli* (Figure S1A). These data show that vimentin enhances the efficiency of phagocytosis of *E. coli* by neutrophils.

Neutrophil extracellular traps (NETs) are large, extracellular, web-like structures composed of cytosolic and granule proteins that are assembled on a scaffold of decondensed chromatin and are released by neutrophils at sites of infection to kill pathogens (43). Vimentin associates with NETs (44), suggesting the presence of vimentin may alter the efficiency of NET formation. To measure NET release, we quantified the percentage of cells that were associated with extracellular DNA. We found that NET release was similiarly efficient in *Vim*^-/-^ neutrophils as compared to the *Vim^+/^*^+^ neutrophils (Figure S1B-C). Our observation indicates that that vimentin does not significantly alter NET formation in these cells.

To examine neutrophil degranulation during LPS stimulation, we measured levels of released matrix metalloprotease 9 (MMP-9). We observed that, at 2h, 4h, and 6h of LPS stimulation, supernatants from *Vim^+/^*^+^ and *Vim^-/-^* neutrophils showed similar levels of MMP-9 by ELISA (Figure S1D). These data show that vimentin has minimal effects on degranulation of neutrophils in our assay.

### Vimentin restricts inflammasome activation and cytokine production in neutrophils

Because vimentin is involved in the inflammatory response (14), we examined the effects of vimentin on cytokine production and inflammasome activity. As TNFα and IL-6 are canonical pro-inflammatory cytokines produced in response to LPS stimulation, we compared the amount of these cytokines released from cells in response to LPS treatment by ELISA. We found that more IL-6 and TNFα were released from *Vim^-/-^* neutrophils at 2h, 4h, and 6h of stimulation with LPS (Figure 2E-F and S1E-F). There were no significant changes in the quantities of IL-6 and TNFα detected from unstimulated cells over time (Figure 2E-F and S1E-F). These results are consistent with previous findings that vimentin restricts inflammatory cytokine production in macrophages (45). In contrast to IL-6 and TNFα, some cytokines such as IL-1β are activated and released from cells by activity of inflammasomes. Previous reports show that assembly of the NLRP3 inflammasome is dependent on the presence of vimentin, and in its absence, IL-1β is not cleaved or released from cells (14). In contrast to previous reports, we observed efficient release of IL-1β from cells lacking vimentin, and by 2 hours of LPS and nigericin stimulation, there was a trend of more IL-1β release (∼1.5 fold) from *Vim^-/-^*neutrophils compared to *Vim^+/^*^+^ neutrophils (Figure 2G). To further test the effects of vimentin on inflammasome activation, we performed western blots to examine activation of caspase-1 in *Vim^-/-^* and *Vim^+/^*^+^ neutrophils under stimulation with LPS and nigericin for 2 hours. Consistently, the levels of pro-caspase-1 were unchanged, but we observed a 1.9-fold increase in total p12/p10 cleavage products in *Vim^-/-^* neutrophils compared to *Vim^+/^*^+^ neutrophils (Figure 2H-J). Together, our data show the inflammasome is activated independently of vimentin in neutrophils.

### Neutrophils lacking vimentin produce more ROS

ROS is an important mechanism for protection against bacteria and serves as first line of defense against pathogens (46, 47). Previous studies have shown that vimentin modulates ROS production in bone marrow-derived macrophages and in the human monocytic cell line THP-1, suggesting it may have similar functions in neutrophils (15). To test whether vimentin deficiency changes neutrophil ROS production, ROS levels were measured by reactivity with cytochrome c (24). We showed that *Vim^-/-^* neutrophils produced 43% more ROS when compared to *Vim^+/^*^+^ cells (Figure 3A). ROS can be produced as byproducts of metabolic processes or can be inducibly produced in response to stimuli through NADPH oxidase complex assembly. When neutrophils were stimulated with PMA to activate PKC, both *Vim^+/^*^+^ and *Vim^-/-^* neutrophils showed similar levels of ROS production and significantly higher levels compared to unstimulated conditions (Figure 3A-B). Because *Vim^-/-^* neutrophils produce more ROS at baseline and there is no difference in the amount of ROS produced upon stimulation, these data suggest that metabolic differences between the cells may be driving the differences in the amount of ROS produced at baseline.

**Figure 3.**
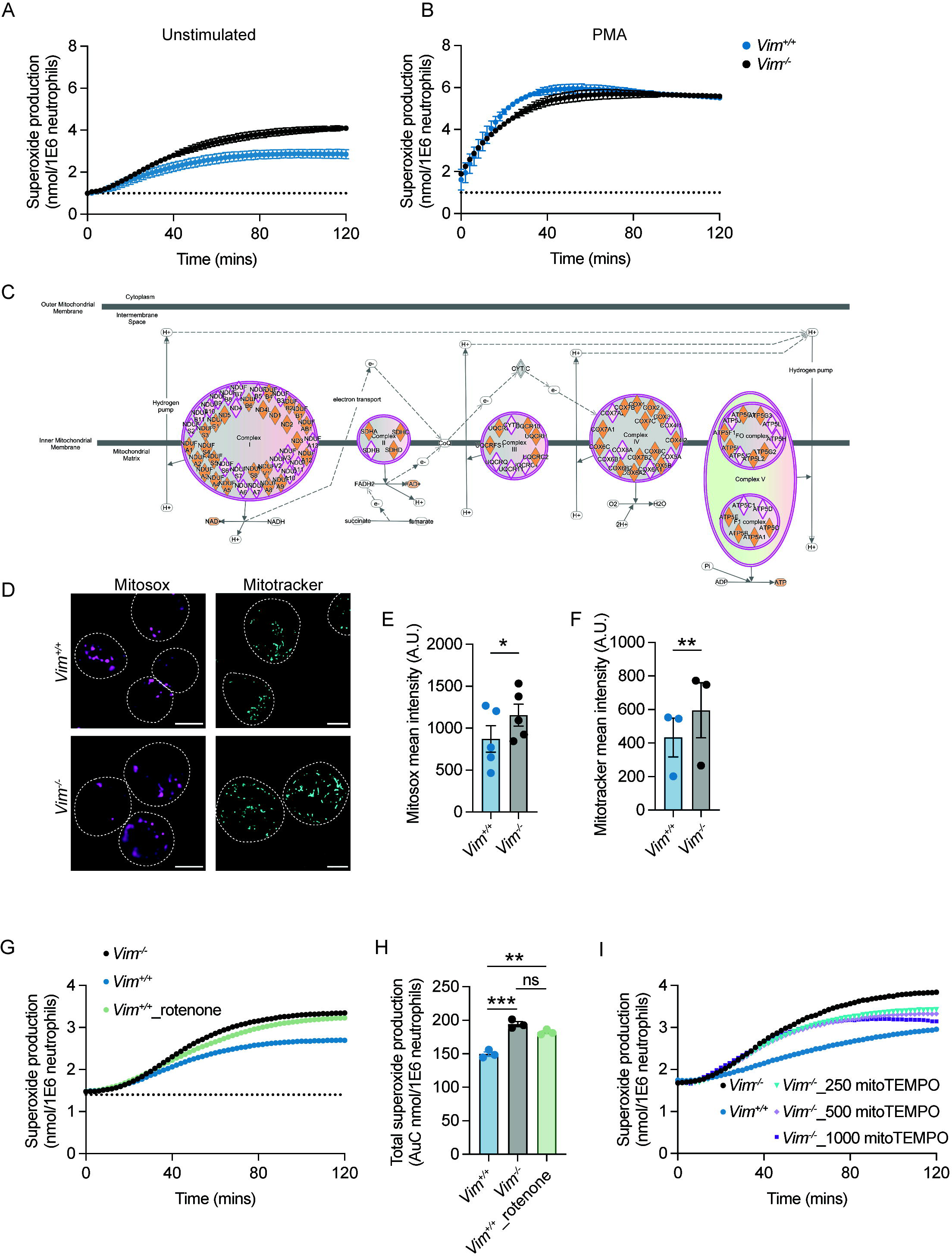
Vimentin regulates mitochondrial ROS production. A-B) ROS production from neutrophils was measured by cytochrome c reactivity in unstimulated cells (A) or PMA-treated cells (B). Blue, *Vim^+/+^*; gray, *Vim^-/-^* neutrophils. C) Ingenuity Pathway Analysis showing multiple genes in electron transport chain complexes are down-regulated in *Vim^-/-^* neutrophils. Pink diamond, downregulated genes in *Vim^-/-^* as compared to *Vim^+/+^*neutrophils. D) Vimentin reduces mitochondria and mitochondrial derived ROS. Representative images. Dotted lines are cells boundaries. Cyan, mitotracker; magenta, mitosox. Scale bar: 5 μm. E) Mitosox intensity per cell (A.U.). F) Mitotracker intensity per cell. G) Inhibitor of complex I of the electron transport chain, rotenone, increases ROS production. H) Quantification of total superoxide production of (G). Dots are independent experiments, and data are mean ± SEM. ns, not-significant; *, p <0.05; **, p<0.01, ***: p-value<0.001. Significance was determined by paired t-test (E-F) or one-way ANOVA with Sidak’s *post hoc* test (H).

### Lack of vimentin leads to mitochondrial dysfunction and mitochondrial ROS overproduction

Previous reports showed that vimentin limits the production of ROS by directly interacting with and inhibiting the formation of the NADPH oxidase complex (15). We observed that ROS production with PMA treatment was unchanged in neutrophils lacking vimentin (Figure 3B), suggesting that, in these cells, vimentin was altering ROS production through a different mechanism. Vimentin loss likely affects multiple pathways and is known to alter the metabolism of fibroblasts by regulating mTOR signaling (48, 49) and is involved in mitochondrial regulation (7, 50–52). Given the importance of ROS production for neutrophil antimicrobial activity, we sought to determine whether analysis of gene expression by RNA-Seq (Figure S2 and Supplemental Material 1) could point to a mechanism for how vimentin affects the functions of neutrophils. To do so, RNA was isolated from neutrophils at baseline and following stimulation with LPS. Given the robust differences in the amount of proinflammatory cytokine produced in response to LPS treatement, we anticipated observing distinct genes activated leading to the production of proinflammatory cytokines such as in the NF-κB signaling pathway, but we observed similar genes activated in response to LPS for both *Vim^-/-^*and *Vim^+/^*^+^ neutrophils (Figure S3). This suggested that the phenotypic difference observed between cells arises not from differential activation of pathways in response to stimulation, but may arise from differences already present at baseline in the cells. Our RNAseq analysis showed 108 genes upregulated and 416 genes downregulated when comparing *Vim^-/-^* and *Vim^+/^*^+^ neutrophils at baseline. To make more sense of how these genes are related, we used our RNAseq data to identify enrichment of genes in particular cellular pathways and to predict the impact of the observed gene expression on pathway activity using Ingenuity Pathway Analysis, and observed significant changes in pathways related to mTOR signaling, EIF2 signaling, and mitochondrial function in *Vim^-/-^* neutrophils compared to the *Vim^+/^*^+^ cells (Figure S4 and Supplemental Material 2). Consistent with our functional data showing PMA stimulation of NADPH oxidase resulted in production of similar amounts of ROS (Figure 3B), there were no changes in NADPH oxidase gene expression in our RNA-Seq data (Supplementary material 1). One mechanism for cells to remove ROS produced via metabolic processes is through the superoxide dismutases SOD1 and SOD2. We observed ∼1.8- and ∼1.7-fold increases in SOD1 and SOD2 in unstimulated *Vim^+/^*^+^ neutrophils as compared to *Vim^-/-^*neutrophils and ∼1.5- and 1.3-fold increases in SOD1 and SOD2 in stimulated *Vim^+/^*^+^ neutrophils compared to stimulated *Vim^-/-^*neutrophils. These data suggest that the greater levels of SOD1 and SOD2 in *Vim^+/+^* neutrophils may lead to lower levels of ROS in these cells. Interestingly, we also noted that there were additional changes to mitochondrial function that are consistent with metabolic changes leading to increased levels of ROS at baseline. Specifically, multiple genes in the electron transport chain (ETC) are downregulated in *Vim^-/-^* neutrophils (Figure 3C) and genes associated with mitochondrial dysfunction were upregulated in the *Vim^-/-^* cells (Supplemental material 2). Together these data suggest that decreased ETC activity in the mitochondria and decreased levels of SOD1/2 may drive the differences observed in ROS levels at baseline in our cells.

To explore whether the changes in ROS production are related to mitochondria, we examined the mitochondrial membrane potential and mitochondrial-derived ROS (mtROS) production by staining *Vim^+/+^* and *Vim^-/-^* neutrophils with mitotracker green and mitosox red, respectively. We observed ∼1.5-fold increase in the intensity of mitotracker and ∼1.3-fold increase in the intensity of mitosox in *Vim^-/-^* neutrophils compared to *Vim^+/+^* neutrophils (Figure 3D-F). These results show that the higher rates of ROS produced by the *Vim^-/-^* neutrophils observed above (Fig. 3A-B) are associated with differences in the amount of ROS produced in mitochondria.

As the expression of ROS scavengers SOD1/2 and multiple genes associated with ETC are downregulated in *Vim^-/-^* compared to *Vim^+/+^*neutrophils, we asked whether we could rescue the ROS levels by manipulating the ETC or SOD activity. To test whether downregulation of ETC genes in *Vim^-/-^* neutrophils drives ROS overproduction in our cells, we inhibited ETC complex I in *Vim^+/+^* neutrophils with rotenone (53). Consistent with our hypothesis, we observed that treatment with rotenone increases the ROS production of *Vim^+/+^*neutrophils by 1.4-fold, which lead to ROS production that was similar to untreated *Vim^-/-^* neutrophils (1.5-fold) (Figure 3G-H). Similarly, to test whether increased levels of the mitochondrial-specific ROS scavengers SOD1/2 contributed to decreased levels of ROS in *Vim^+/+^* neutrophils, we increased the levels of mitochondrial specific ROS scavengers in *Vim^-/-^* neutrophils with mitoTEMPO, a SOD mimetic that functions specifically in the mitochondria. We found that by increasing the amount of mitochondrial-specific ROS scavenger with mitoTEMPO, there was a 0.84-fold reduction of ROS production in *Vim^-/-^* neutrophils (Figure 3I). These data show that decreases in ETC and SOD1/2 drive ROS overproduction in *Vim^-/-^*neutrophils.

## Discussion

Vimentin is important for regulating neutrophil function. Here, we showed that the absence of vimentin in neutrophils caused defects in swarming and phagocytosis, but not NETosis and degranulation (Figure 2A-D and S1B-D). We observed that loss of vimentin increased TNFα, IL-6, IL-1β, and inflammasome activation (Figure 2E-J). We also showed a significant increase in mitochondrial ROS production and dysregulation of mitochondrial genes in *Vim^-/-^* compared to *Vim^+/+^* neutrophils using mitosox, mitotracker, and RNA-Seq data (Figure 3A-F). We demonstrated that we can alter changes in ROS production associated with vimentin by inhibiting mitochondrial complex I or using a SOD mimetic (Figure 3G-H). Altogether, these data suggest seemingly opposing roles for vimentin in regulating the antimicrobial activity of neutrophils (Figure 4).

**Figure 4:**
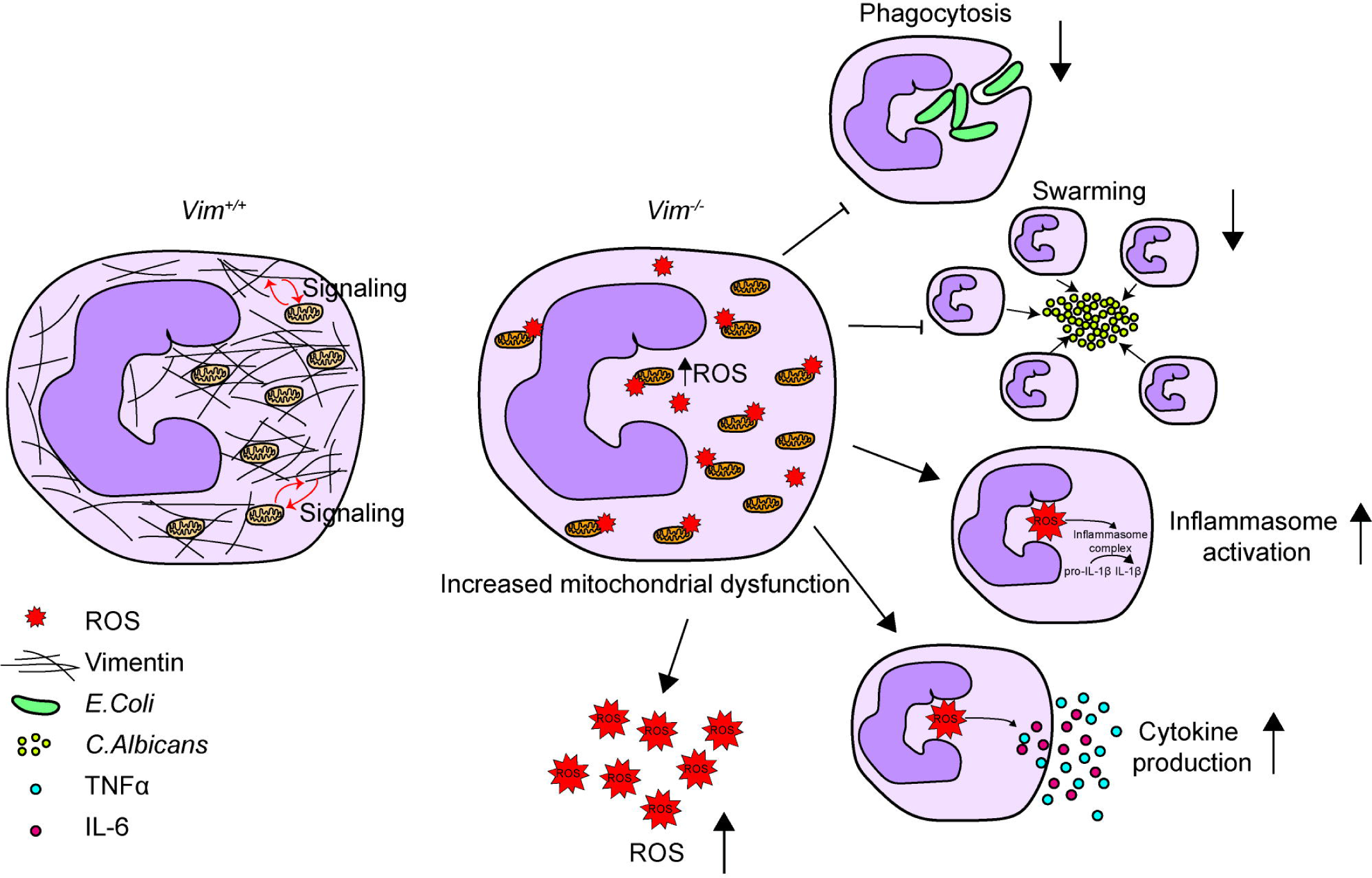
Schematic showing vimentin function in neutrophils. Vimentin supports mitochondrial function. In the absence of vimentin, mitochondria become dysregulated and produce more ROS. Vimentin loss also leads to decreased phagocytosis and swarming and increased inflammasome activation and proinflammatory cytokine production.

Phagocytosis, swarming, and NETosis are mechanisms neutrophils require to clear pathogens (54). Whereas these functions can be induced by cytokines and ROS (54–57), our data support a more direct requirement for vimentin in the ability of neutrophils to traffic to pathogens. We observed defects in swarming and phagocytosis despite increased production of IL-6, IL-1β, and ROS from *Vim^-/-^* neutrophils, which is consistent with previous reports showing defects in motility in cells lacking vimentin (10, 21, 22, 39, 40).

Our results suggest a distinct mechanism by which vimentin affects inflammatory activation of neutrophils through regulation of mitochondrial ROS production. Whereas previous studies showed that differences in ROS production in macrophages lacking vimentin is derived from the NADPH oxidase complex (15), our studies show mitochondrial dysfunction leads to increased levels of ROS in neutrophils lacking vimentin. In neutrophils, mitochondria lack life-supporting activity but regulate cell death and ROS production (58). Previous studies have shown that neutrophils express an extremely low amount of cytochrome c, which likely leads to accumulation of electrons in the respiratory chain and subsequent formation of ROS (58). Here, we showed that *Vim^-/-^* neutrophils produce more mitochondria-specific ROS than *Vim^+/+^* neutrophils (Figure 3A, C-F).

Previously, vimentin was shown to be required for NLRP3 activation in macrophages (14). In contrast, we observe that vimentin is dispensible for NLRP3 activation in neutrophils. NLRP3 is regulated in neutrophils in a distinct manner from macrophages such that the components of the inflammasome are present in vacuoles (59), which suggests that the distinct regulatory mechanisms may be independent of vimentin in these cells and/or alternative scaffolds may be used for the assembly of the inflammasome. Alternatively, mitochondria-derived ROS can lead to the activation of the NLRP3 inflammasome (60, 61), suggesting that the increased levels of michondiral-derived ROS may overcome the requirement of vimentin in these cells. Yet, macrophages produce more ROS through the NADPH oxidase complex upon vimentin knockout (15), which would suggest that ROS in general is unable to overcome vimentin-dependent inflammasome assembly and that, if the ROS was activating the inflammasome in our system, the ROS would likely either need to be derived from specific pools of ROS produced by mitochondria or related to the spatial location of the ROS produced by the mitochondria in these cells.

Together, our study suggests that vimentin regulates neutrophil functions by 1) supporting cellular structure and cell motility and 2) regulating mitochondrial ROS production. However, the mechanism of how vimentin regulates mitochondrial function is unknown and requires further investigation. As neutrophils are commonly involved in several chronic inflammatory diseases and infection, a number of studies have focused on targeting neutrophil function as a possible treatment. Thus, vimentin could be used as a target for manipulating neutrophil functions.

## Supporting information

RNA Sequencing data

Pathway Analysis

## Data Availability Statement

Further information and requests for reagents should be directed to and will be fulfilled by Brian Russo.

## Author Contributions

B.C.R., J.T., T.N.H., A.S., M.K.M., and D.M designed the experiments. T.N.H., J.T., A.S., A.G., T.B., and B.C.R. performed the experiments. T.N.H., J.T., and B.C.R. wrote the manuscript and all authors discussed the results and commented on the manuscript.

## Acknowledgements

We would like to thank Arienne Theiss, Marijke Keestra-Gounder, Kelly Doran for reagents, Kyla Ost for *C.albicans*, and Alex Hopke and Daniel Irimia for the printed micro scale swarming array slides. We would like to thank Marcia Goldberg, Peter Hensen, Mercedes Rincon, Leslie Berg, Cristina Penaranda, Ross Kedl, Jared Klarquist, Marijke Keestra-Gounder, and the Keestra-Gounder lab for helpful discussions. This work was partly funded by K22 AI137296, R35 GM146923, and funds of the University of Colorado and the University of Michigan to B.C.R., and by R01 AI132638 to M.K.M..

