## Supplementary Figures for "Vimentin regulates mitochondrial ROS production and inflammatory responses of neutrophils"

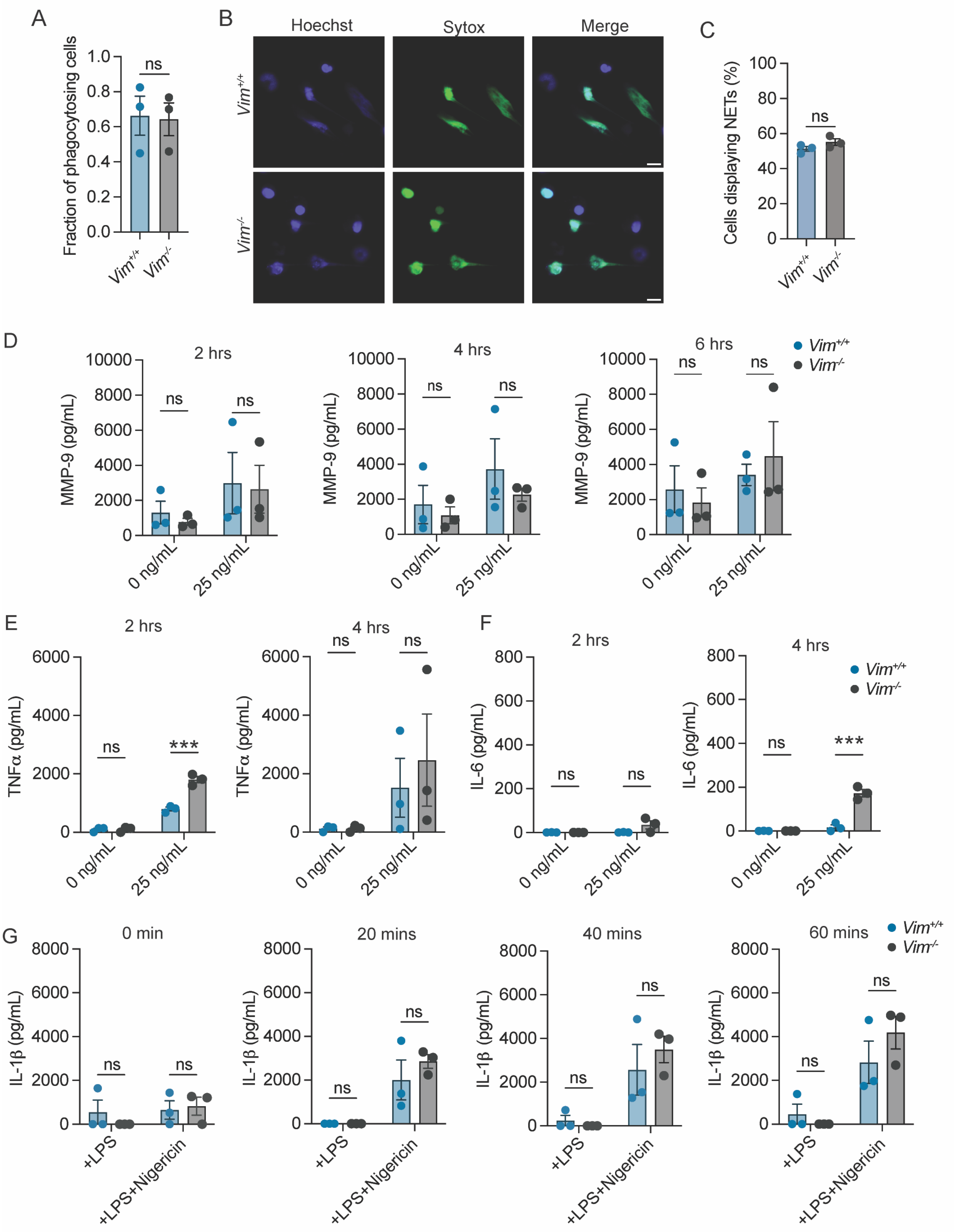

**Figure S1. Neutrophil functions that are not affected by vimentin.**

A) Fraction of phagocytosing cells. B) Representative images of NET formation induced by A23187. Green, extracellular DNA; blue, all DNA. Scale bar: 10 $\mu$m. C) Quantification of percentage of cells displaying NETs in $Vim^{+/+}$ and $Vim^{-/-}$ neutrophils from images represented in B. D) Levels of MMP-9 (pg/mL) released from neutrophils treated with LPS (0 and 25 ng/mL) at 2, 4, and 6 hours by ELISA. E-F) Quantification of TNF$\alpha$ (E) or IL-6 (F) produced by $Vim^{+/+}$ and $Vim^{-/-}$ neutrophils in response to LPS stimulation at 2 and 4 hours. G) Quantification of IL-1$\beta$ produced by $Vim^{+/+}$ and $Vim^{-/-}$ neutrophils in response to treatment with LPS alone or LPS and nigericin at 0, 20, 40, and 60 minutes. Data are mean ± SEM from three independent experiments with 5 $Vim^{+/+}$ or 6 $Vim^{-/-}$ cell lines. Dots are independent experiments. ns, not significant by paired t-test (A, C) or two-way ANOVA with Sidak's *post hoc* test (D-G).

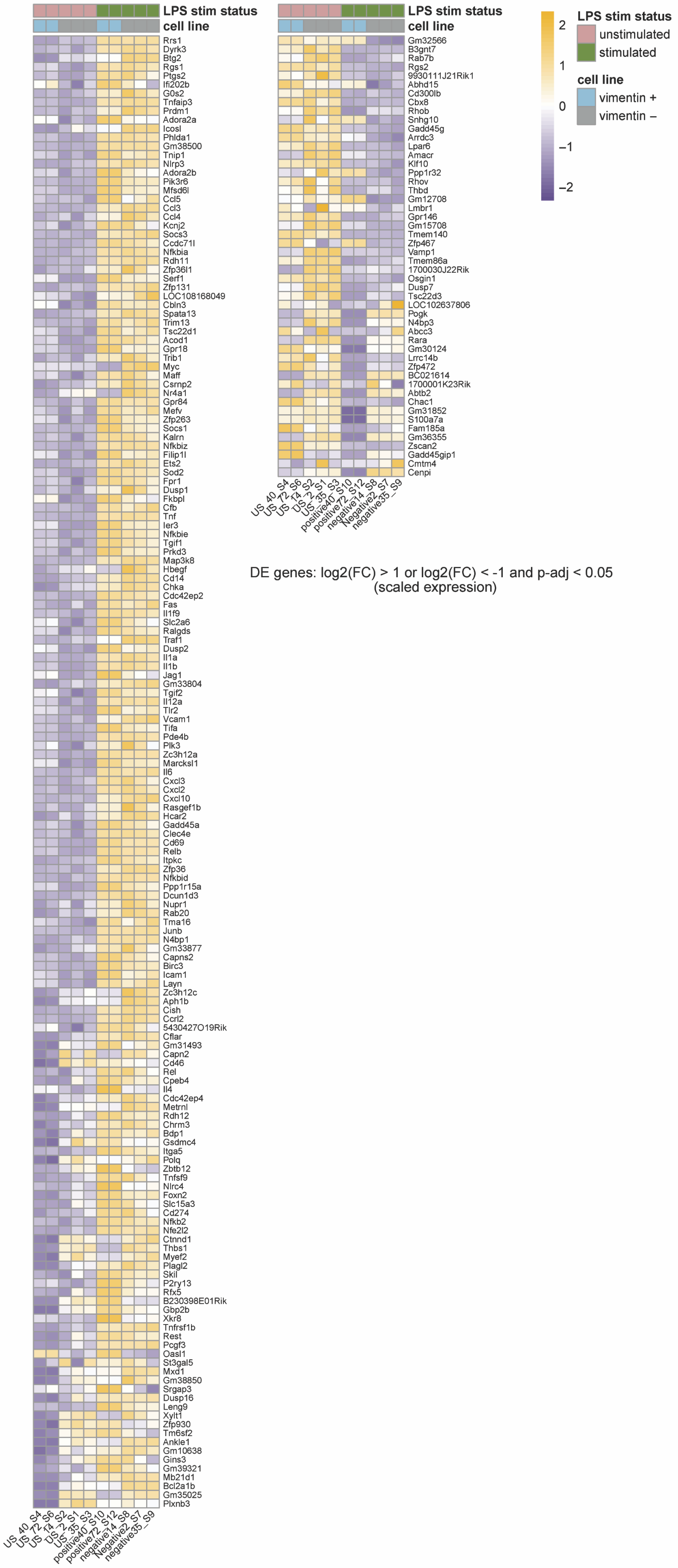


DE genes: log2(FC) > 1 or log2(FC) < -1 and p-adj < 0.05
(scaled expression)

**Figure S2. Heatmap of gene differential expression.**

Heatmap showing differential expression genes with $\log_2(\text{fold\_change}) > 1$ or $\log_2(\text{fold\_change}) < -1$ with adjusted p-value < 0.05 in all conditions.

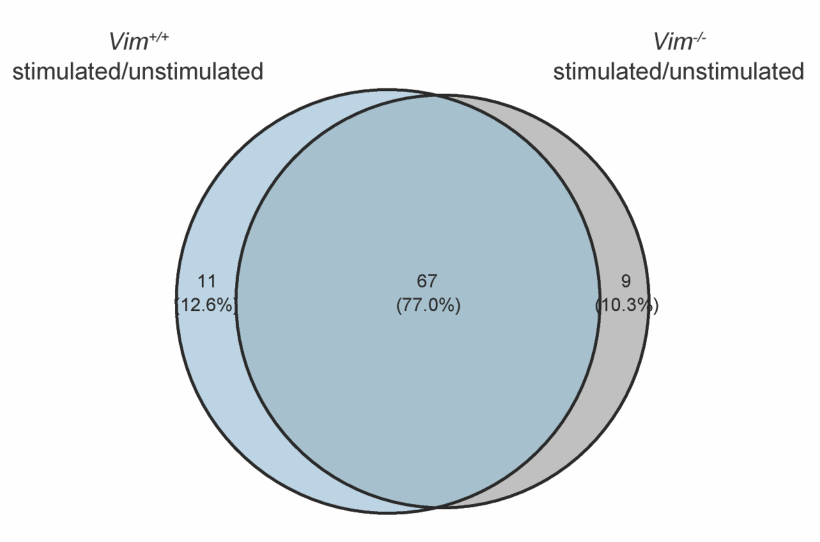


**Figure S3. LPS stimulation affects similar genes between *Vim*$^{+/+}$ and *Vim*$^{-/-}$ for TNFα signaling via NF-κB hallmark.**

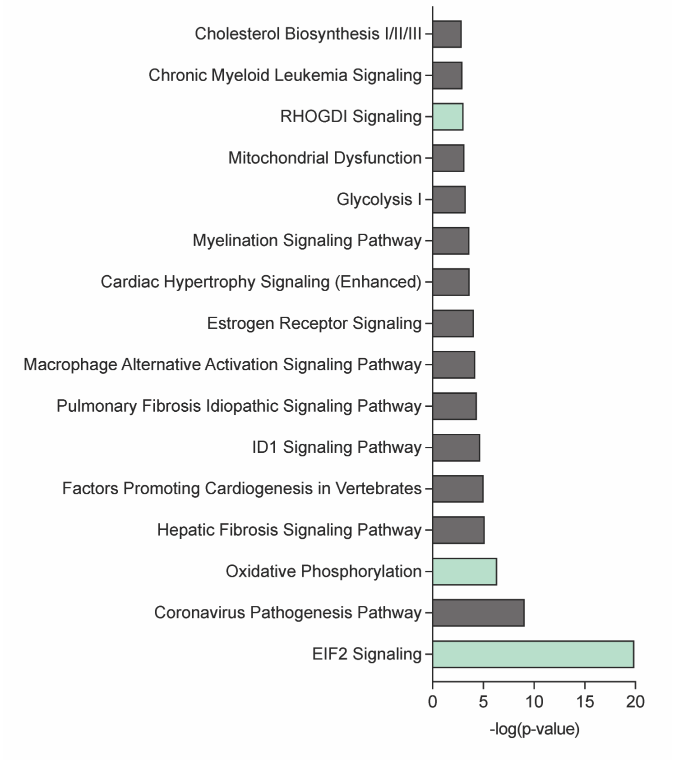


**Figure S4. Bar graph showing pathways altered in *Vim*$^{-/-}$ compared to *Vim*$^{+/+}$ neutrophils.** The top 20 pathways affected with the highest -log(p-value) from Ingenuity Pathway Analysis were selected and the pathways with the z-score > 2 or z-score < -2 were considered significant. Gray, upregulated in *Vim*$^{-/-}$ cells; green, downregulated in *Vim*$^{-/-}$ cells.

**Supplemental Material 1. Raw RNA-Seq data.**

**Supplemental Material 2. Pathways analysis data from Ingenuity Pathway Analysis.**
